# Segmentation and Analysis of Anterior Lamina Cribrosa Surface using Non Local MRF and Metropolis Hasting Algorithm

**DOI:** 10.1101/2020.05.30.125682

**Authors:** Abhisha Mano

## Abstract

The segmentation of anterior Lamina Cribrosa surface from the OCT image is an essential task for analysis of glaucomatous damage. A Bayesian method is used to segment LC surface whereas prior knowledge about shape and position of LC layer is obtained by the non local Markov Random field and K-means segmentation. The Metropolis-Hastings (MH) algorithm provides autocorrelation graph and distribution of samples from a probability distribution. By using this technique acceptance probability is calculated. Finally, the LC layer is analysed whether it is normal or abnormal. This technique provides an accuracy of 96.7%

## 1. Introduction

The Lamina Cribrosa (LC) is used in the identification of glaucomatous injury. The Lamina Cribrosa (LC) is a eye layer which gives rise to retinal ganglion cell that extends to the brain with visual data. A boundary is formed by the Lamina Cribrosa in between the intraocular space and retrobulbar space with intraocular pressure and retrolaminar pressure respectively. In optical coherence tomography (OCT) optic nerve head ONH structures can be viewed clearly. Nowadays, spectral domain SD-OCT Enhanced depth imaging (EDI) was used to view the retinal structures. The first and final layer of Lamina Cribrosa are inner limiting membrane and retinal pigment epithelium respectively. The internal limiting membrane is used to diagnose various interface diseases, in the macular area. The pigmented layer nourishes retinal visual cells. The pressure of eye is measured in millimeters of mercury (mm Hg). The normal eye has a pressure of 12-22 mm Hg. Eventhough the pressure is greater than normal range, and there is no symptoms of glaucoma it is called as ocular hypertension. If pressure is between 12mm Hg to 22 mm Hg then the Lamina Cribrosa layer is in normal stage else it is in abnormal stage.

In method[1],automatic segmentation of slices of LC are done in 3D OCT imaging.BM4D algorithm performs grouping and filtering. A 4D transform will exploit the nonlocal correlation occurring among the pixels. By inverse transformation estimates are obtained for grouped data[8].Otsu threshold method is used to identify the glaucoma from the disease[11].By artificial neural network, optic disc size and rim area is estimated[4].Here bruch’s membrane is located using a deconvolution method. By solving MAP-MRF labeling problem, contour is formed, which is used in segmentation [5]. Many LC segmentation techniques [2],[3].[6],[7],[9],[10]and the structures are analysed [12][13][14][15][16][17] and explained. An approach for segmenting the retinal images using superpixel is proposed in paper[18]. An wavelet based segmentation method is applied for DNA fragment sizing in AFM images and for mammogram images [19][23]. An encryption techniques in images using mutation is presented in paper[21].An optimization based segmentation of brain images is presented in paper[20].Edge detection of microarray images by applying wavelet technique is presented in paper [22].

This paper explains a method for segmentation of Lamina Cribrosa surface. A surface evolution method with shape parameter is used. The shape is varied by Markov random field in an iterative manner. The parameters are computed using a Metropolis-Hastings algorithm and the surface is analysed whether it is normal or abnormal.

## 2. Proposed Method

Optical coherence tomography (OCT) is taken as input and given to the layer segmentation step. The first and the final layer of the Lamina Cribrosa, inner limiting membrane and retinal pigment epithelium respectively are segmented in this step. Then preprocessing is done where the area of focus is determined. A Bayesian method is used to segment LC surface whose prior knowledge about shape and position of LC layer is obtained by the non local Markov Random field and K-means segmentation.

### 2.1. Non Local Markov Random Field Segmentation

In non-local range MRF, the clique of each and every pixel has local neighbourhood patch whose topmost patches is obtained by block matching. The non local range of pixels are obtained from the image region based on similar patches. The filter that is applied over the non-local range is a 3-dimensional filter. Markov Random Field (MRF) will remove the correlation of intensities among voxels. The non local approach will remove the repeated structures in an image. The gradients of loss function are computed. Hence optimization based on gradients can be applied.

### 2.2. Metropolis Hasting Algorithm

Metropolis-Hastings algorithm a the sampling method is inspired by the Biased and Filtered Point Sampling (BFPS) method.

#### Algorithm

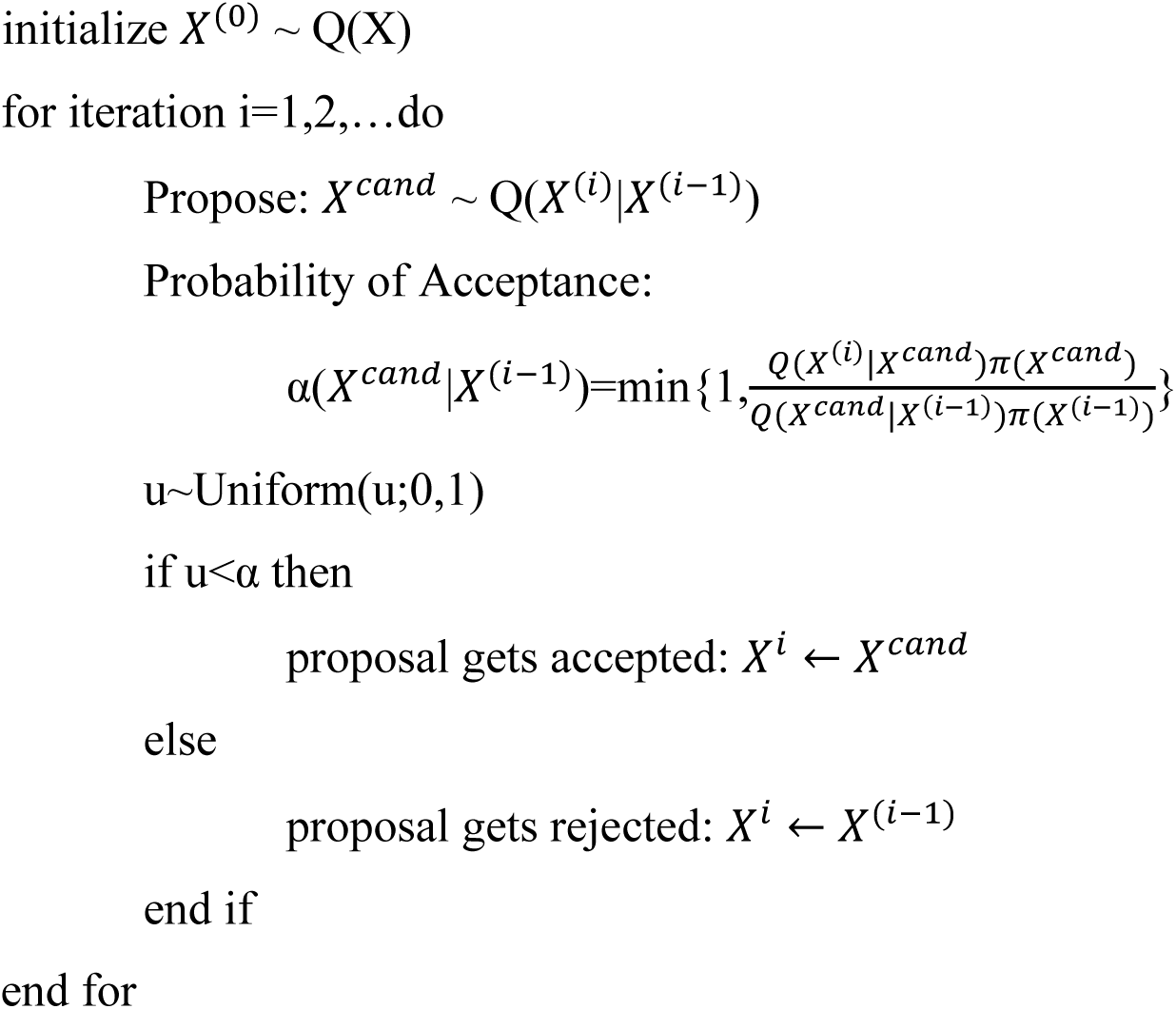

First initialize the samples. Generate a proposal and compute the acceptance probability, followed by accepting the candidate sample having a specified probability.

## 3. Implementation

Optical coherence tomography (OCT) image is taken and given as input to the system. Fig 2 shows the OCT image

**Fig: 1.**
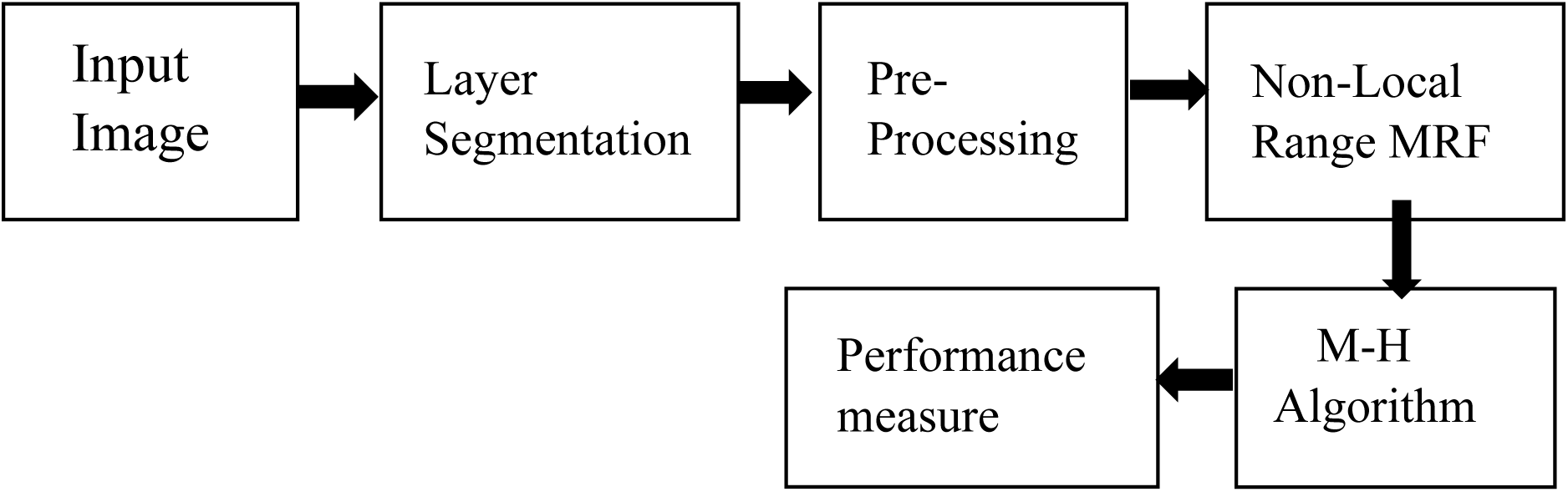
Block Diagram of Proposed Method

**Fig 2.**
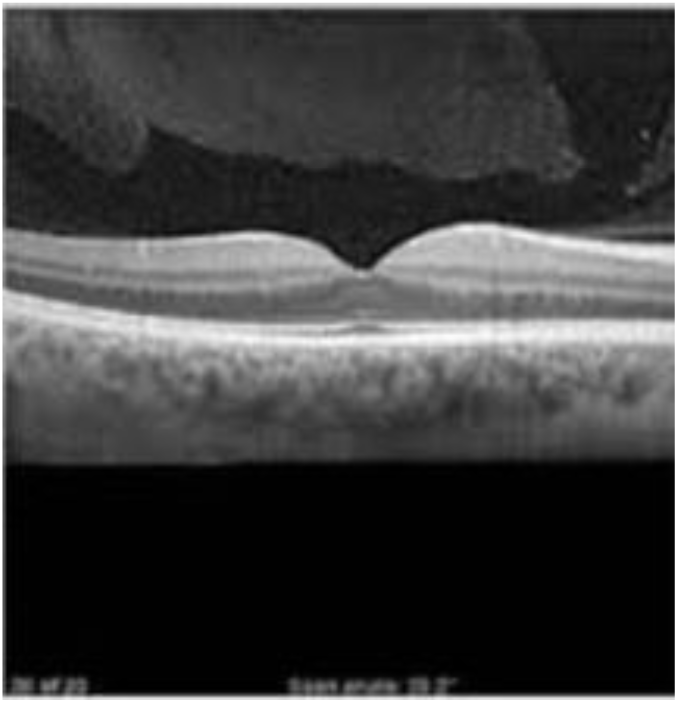
Input Image

The first layer of the Lamina Cribrosa i.e, inner limiting membrane and the final layer of the Lamina Cribrosa i.e, retinal pigment epithelium is segmented by the layer segmentation process. Fig 3 shows the segmented layers. The boundary of the Lamina Cribrosa layer is segmented using canny edge detection and the blurred image is obtain by using Gaussian blur. Fig4a and Fig 4b shows the boundary of image and blurred image respectively.

**Fig 3.**
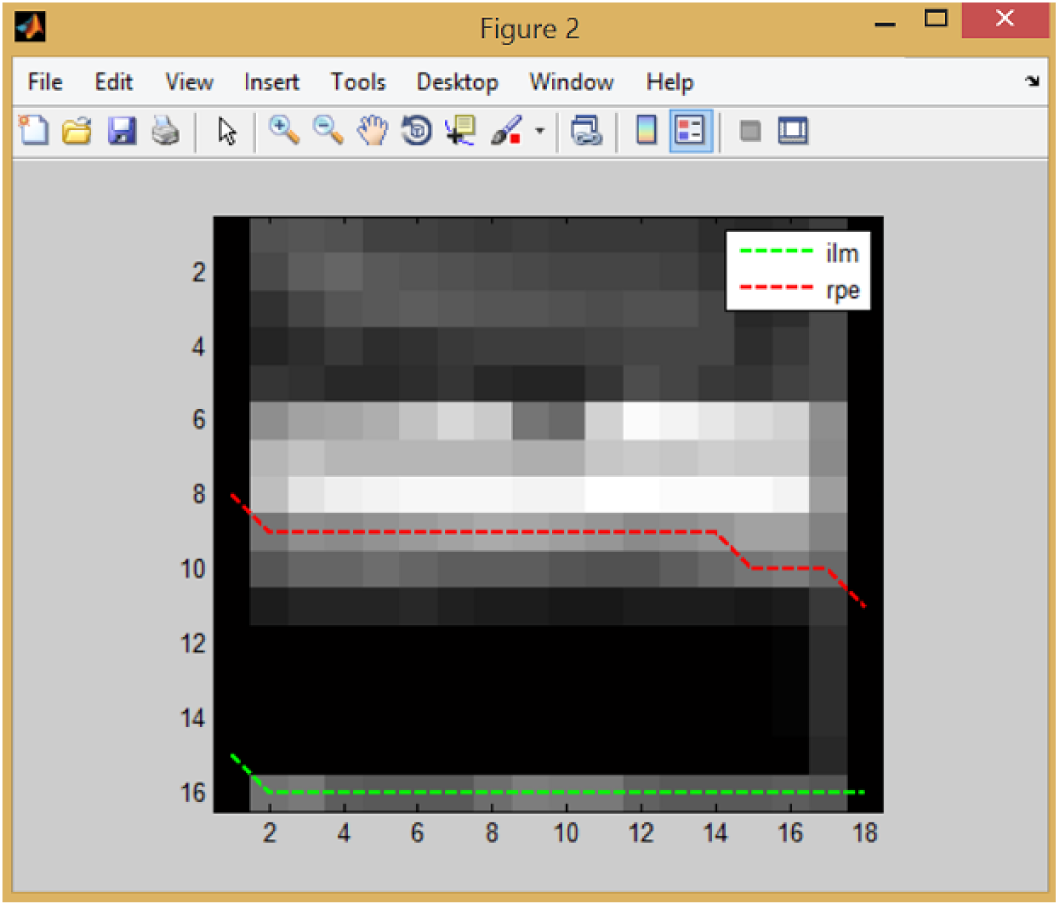
Segmented Layer

**Fig 4a.**
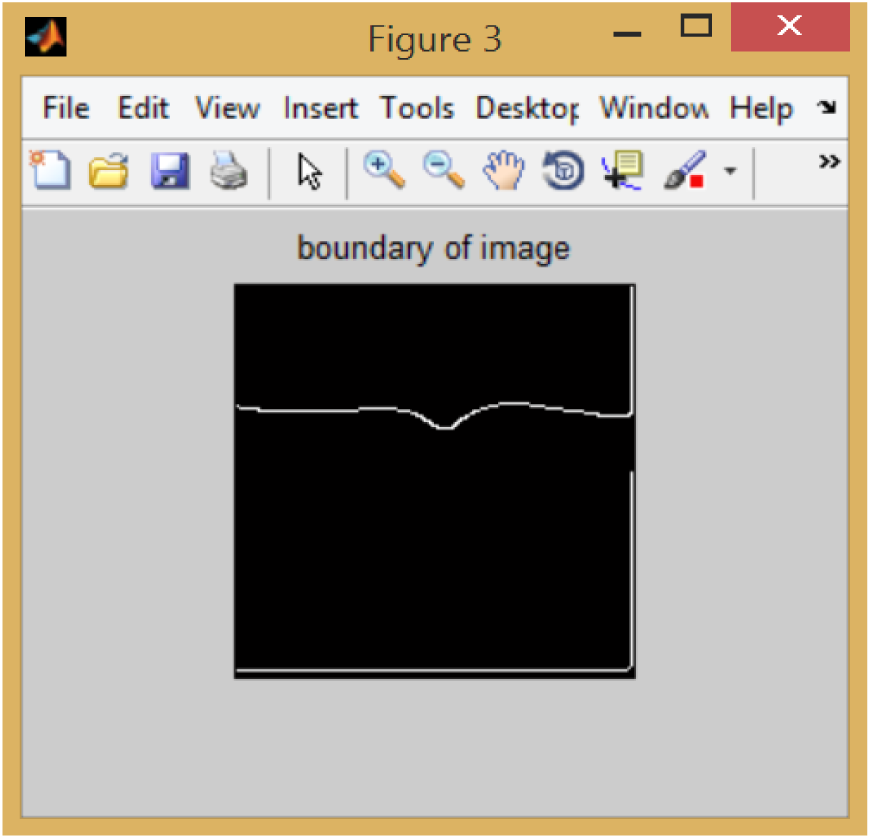
Boundary of the Image

**Fig 4b.**
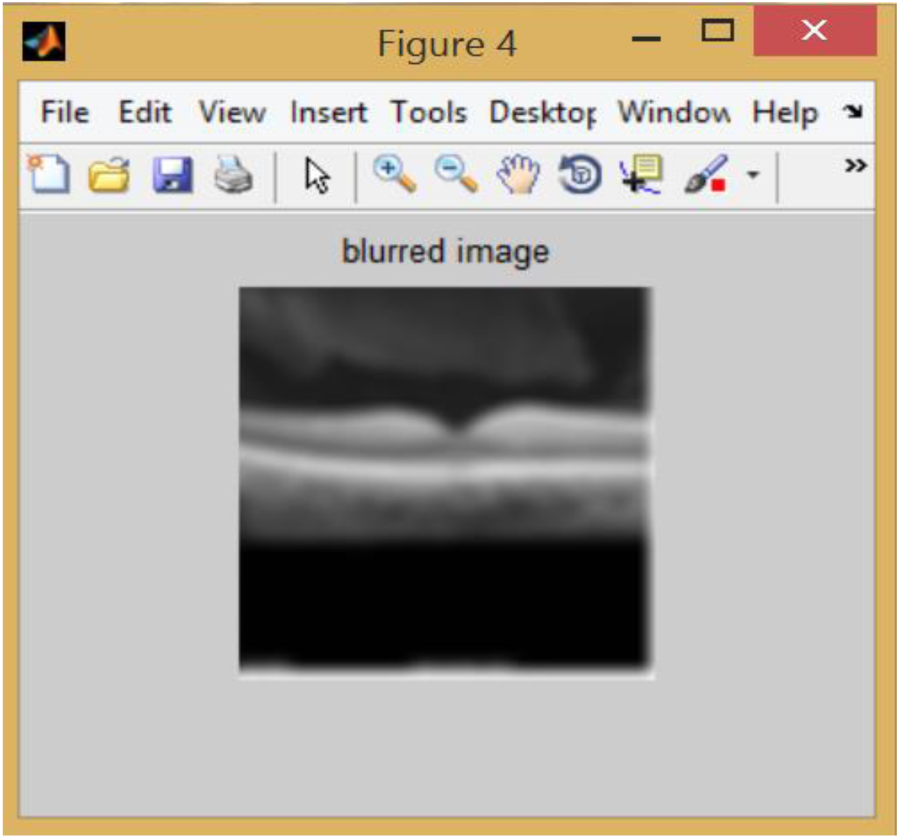
Blurred Image

From the prior knowledge of K-means segmentation initial label is obtained from which the final label can be obtained using Non local Markov random field segmentation. Figure 5 shows the initial label segmented output.

**Fig 5.**
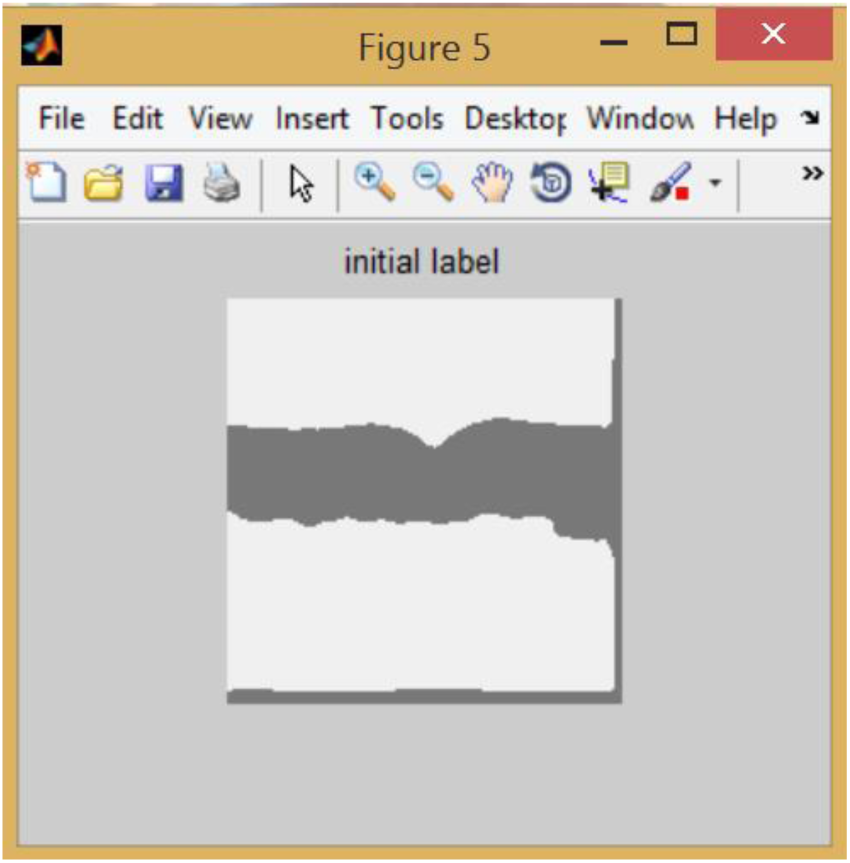
K-means segmented output

Fig 6 shows the final label of the segmented output.

**Fig 6.**
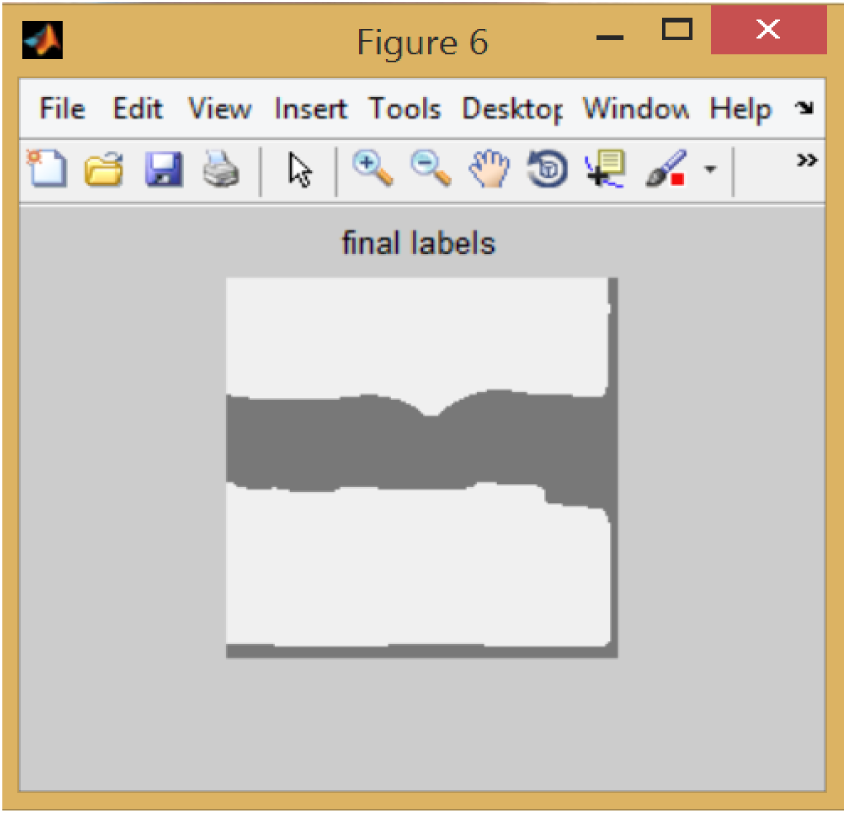
NLMRF segmented output

With the help of the initial and final labels, the density value of the Lamina Cribrosa is found. The density graph is shown in the Fig 7. The graph is plot between the EM iteration and the sum of U. Sum of U denotes the value of initial labels and final labels which is obtained from the segmentation process. The value of EM iteration given as input is 10.

**Fig 7.**
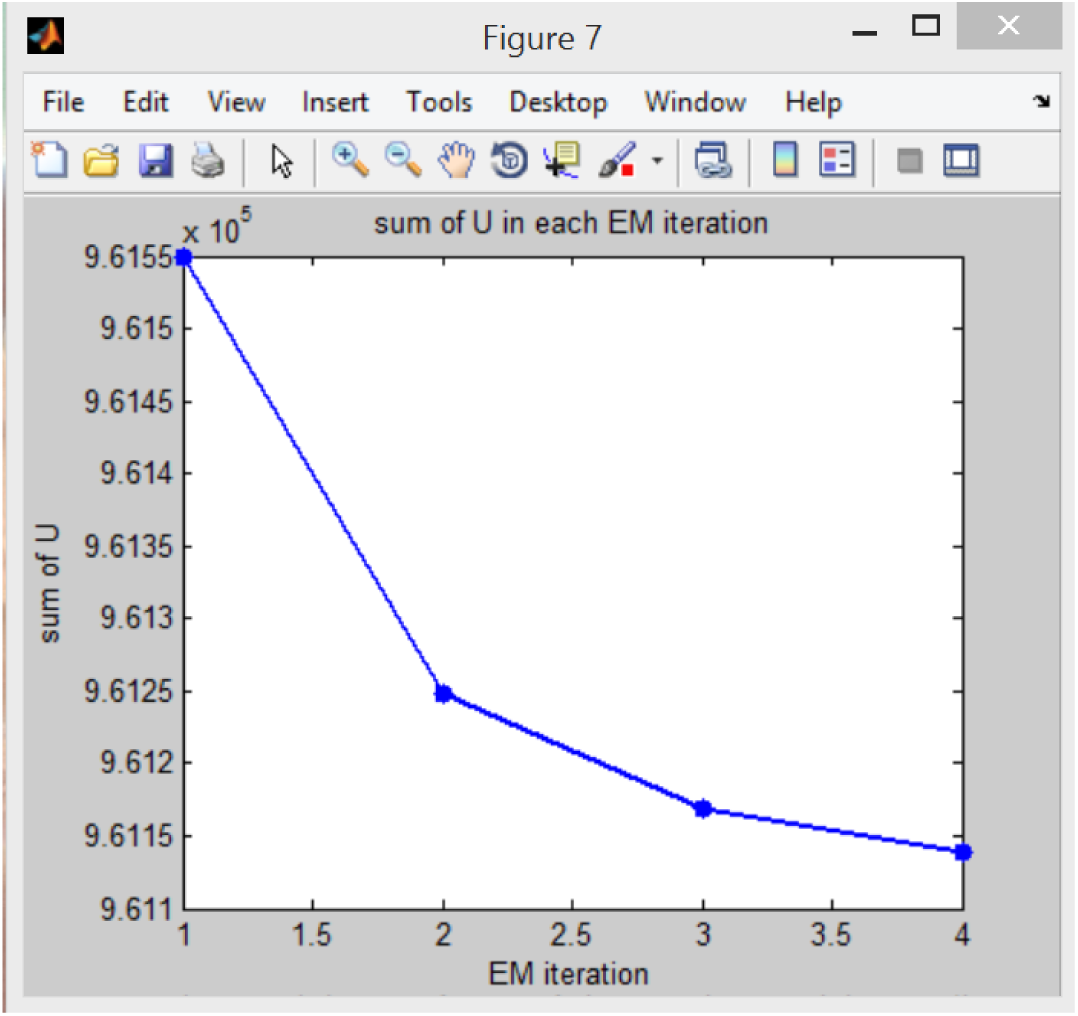
Density Graph

The Metropolis-Hastings (MH) algorithm provides autocorrelation graph and distribution of samples from a probability distribution. Here we choose 10000 samples for the process. The autocorrelation graph shows the values for the first 100 samples as well as last 100 samples.Fig 8 shows the autocorrelation graph obtained.

**Fig 8.**
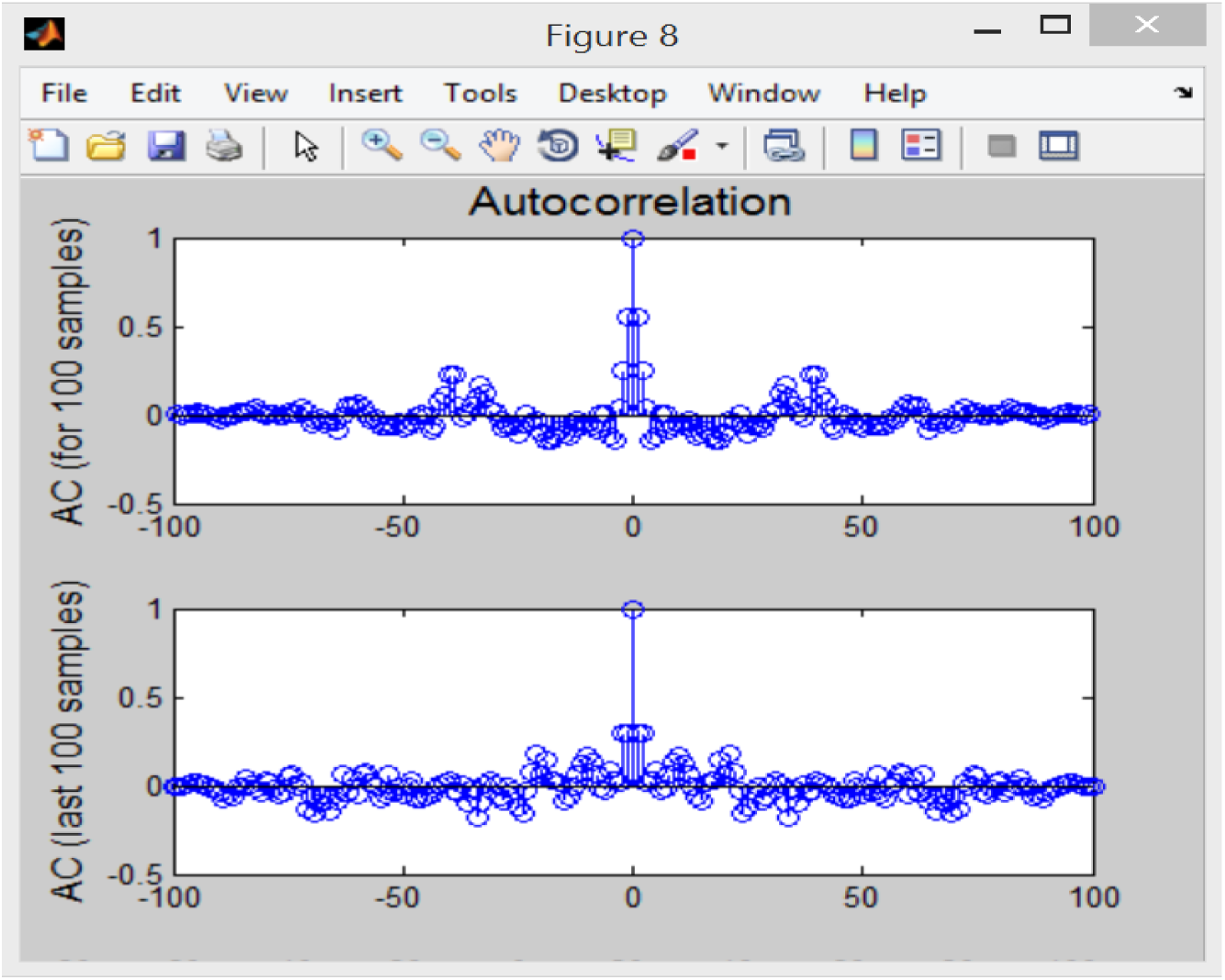
Autocorrelation Graph

Fig 9 shows the distribution of samples graph. This graph is drawn with the help of probability density function. It provides the value of acceptance rate as 0.5418.

**Fig 9.**
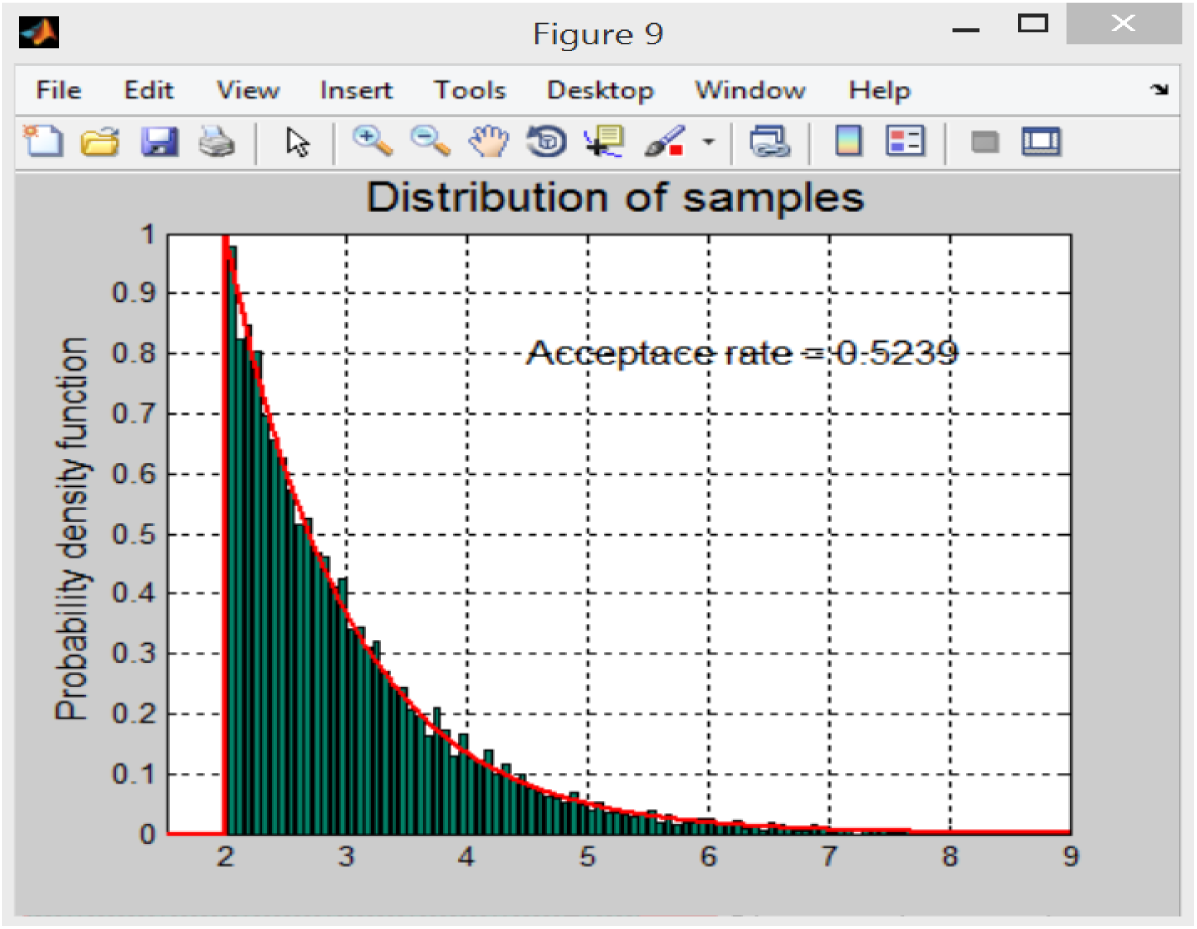
Distribution of sample graph

With the help of the density, autocorrelation and the distribution of values the Lamina Cribrosa is verified whether it is in normal stage or in abnormal stage. If the value is below 12 mm Hg then the help box will show as low pressure. If the value is above 22 mm Hg then the help box will show as high pressure.

The value of pressure obtained from this image is 31.8256. Thus the help box shows as high pressure.

## 4. Conclusion

The work proposed in this paper is a shape based surface evolution method for the segmentation of the anterior LC. A non local approach gives a MRF energy function. Metropolis-Hastings (MH) algorithm provides the accurate density, autocorrelation and distribution of values. The accurate value of pressure is obtained from which the Lamina Cribrosa is analysed whether it is normal or abnormal. Accuracy of this approach is 96.7%

